# Roots fuel cell produces and stores clean energy

**DOI:** 10.1101/2022.09.01.506267

**Authors:** Yaniv Shlosberg, Ailun Huang, Tünde N. Tóth, Richard B. Kaner

**Affiliations:** Department of Chemistry and Biochemistry, University of California at Santa Barbara, Santa Barbara, CA, 93106, United States; Center for Polymers and Organic Solids, University of California at Santa Barbara, Santa Barbara, CA 93106, United States; Department of Chemistry and Biochemistry, Department of Materials Science and Engineering, and California NanoSystems Institute, University of California, Los Angeles, CA, USA; Biological Research Centre, Hungarian Academy of Sciences, Szeged, Hungary

## Abstract

In recent years, extensive scientific efforts have been conducted to develop clean bio-energy technologies. A promising approach that has been under development for more than a hundred years is the microbial fuel cell (MFC) which utilizes exo-electrogenic bacteria as an electron source in a bio-electrochemical cell. The viability of bacteria in soil MFCs can be maintained by integrating plant roots which release organic materials that feed the bacteria. In this work, we show that rather than organic compounds, roots also release redox species that can produce electricity in a bio-fuel cell. We first study the reduction of the electron acceptor Cytochrome C by green onion roots. We integrate green onion roots into a bio-fuel cell to produce a continuous bias-free electric current for more than 24 h in the dark. This current is enhanced upon irradiation of light on the onion’s leaves. We apply cyclic voltammetry and 2D-fluorescence measurements to show that NADH and NADPH act as major electron mediators between the roots and the anode, while their concentration in the external root matrix is increased upon irradiation of the leaves. Finally, we show that roots can contribute to energy storage by charging a supercapacitor.

## 1. Introduction

In recent years, many efforts have been made to replace fossil fuels with green energy technologies. The utilization of MFCs for electricity generation has been extensively studied for more than a hundred years.(Bond and Lovley, 2003; Menicucci et al., 2006; Min et al., 2005; Rabaey et al., 2005; Ramu et al., 2020; Rhoads et al., 2005; Ringeisen et al., 2006; Slate et al., 2019; Wei et al., 2016) In this method, bacteria acts as an electron donor at the anode (Fang et al., 2020) or an electron acceptor at the cathode (Bergel et al., 2005; Fang et al., 2020). The electron transfer between the exo-electrogenic bacteria and the anode may be conducted by either of two mechanisms: mediated electron transfer (MET) (Bond et al., 2002; Bond and Lovley, 2003; Chaudhuri and Lovley, 2003; Liu et al., 2006; Min et al., 2005; Pham et al., 2003) or direct electron transfer (DET) (Sekar et al., 2016; Thirumurthy et al., 2020; Wei et al., 2016). In the case of MET, redox-active metabolites such as derivatives of flavins, phenazines, or NADH (Shlosberg et al., 2023a) are secreted from the cells to reduce the anode. The MET can be enhanced by the addition of an exogenous electron mediator such as potassium ferricyanide (Huang et al., 2015; Lan et al., 2013; Laohavisit et al., 2015; McCormick, 2013, 2011; Ochiai et al., 1983). DET is performed by conductive *pili* (Heidary et al., 2020; Lovley, 2012; Nevin et al., 2008; Yi et al., 2009) or metal respiratory complexes (Hartshorne et al., 2007; Shi et al., 2012). Among the species that perform a high degree of DET are *Shewanella onedis* and *Geobacter* (Hartshorne et al., 2007; Heidary et al., 2020; Lovley, 2012; Nevin et al., 2008; Shi et al., 2012; Yi et al., 2009). The power generation of MFCs can be enhanced by implementing them into the soil, while environments with higher organic carbon content and lower pH enhance the power generation (Jiang et al., 2016).

Another approach that evolved from MFCs is the utilization of live photosynthetic unicellular organisms such as cyanobacteria or microalgae (Grattieri, 2020; Grattieri et al., 2019a, 2019b; Shlosberg et al., 2021b, 2020; Tschörtner et al., 2019). In this case, the electron source is photosynthesis where the electron mediation is mainly conducted by the redox couple NADP^+^/NADPH that can cycle between the photosystems inside the organisms and the external anode (Shlosberg et al., 2020). The photocurrent can be enhanced and prolonged by the addition of exogenous NAD^+^, NADP^+^, or vitamin B1 (Shlosberg et al., 2022f, 2021b, 2020) that can enter the cells and accept an electron from photosystem I. Exogenous artificial electron mediators such as potassium ferricyanide (Lee et al., 2020) can also enhance the photocurrent by accepting electrons at the external surface of the cells. Recently, it was discovered that bio-photo electrochemical cells are not limited to unicellular organisms and can be directly produced from macroalgae (seaweeds) (Shlosberg et al., 2022b) and green tissues of terrestrial plants such as leaves and stems (Shlosberg et al., 2022c). The photocurrent production of such systems is about 2-3 orders of magnitude greater than for microorganism-based bio-photo electrochemical cells (BPECs) (Shlosberg et al., 2022d). Interestingly, photocurrent may also be harvested from non-photosynthetic cells and organisms such as non-photosynthetic bacteria, (Shlosberg et al., 2023a) and fruits (Shlosberg et al., 2023b).

Rather than using live organisms, photoelectricity generation can be accomplished by isolating photosynthetic components at different degrees of purification, such as chloroplasts, Thylakoid membranes, and photosystems (Efrati, 2016; Efrati et al., 2013; Hartmann et al., 2020; Hasan et al., 2017; Li, 2016; Yehezkeli et al., 2012; Zhao et al., 2014) or by catalysis of isolated enzymes (Herkendell, 2021; Herkendell et al., 2017). In contrast with live organisms in which the addition of an external mediator is optional, isolated compounds are incapable of performing EET by themselves, and require the addition of an exogenous electron mediator (Pinhassi et al., 2016). Rather than making a specific isolation of a component of interest, a different approach utilizes yeast extracts. Similar to bacteria, live yeast may also conduct exo-electrogenic activity in MFCs (Bahartan et al., 2012). Yet, most of the reducing molecules are located inside the cell cytoplasm. Luria-Bertani media which is mostly based on yeast extract consists of a rich content of components which can produce photocurrent in biofuel cells (Shlosberg et al., 2022e).

An additional interesting method is plant-microbial fuel cells which integrate soil MFCs with plant farming. In this approach, plants release through their roots organic materials that are consumed by the bacteria enabling their cultivation in the soil (Moqsud et al., 2017; Strik et al., 2008).

Rather than secreting components, roots can sense and electrochemically interact with their environment. Such a phenotype can be observed in the electro-tropism responses of the roots. Upon application of an electric field, roots can decompose the starch granules that direct them toward gravitropism (elongation along the gravitational direction) and re-orient their growth direction (Cassab, 2007; Volkov et al., 2013). The electro-tropic mechanism involves biomolecules that are capable of charge transfer. Integration of these electro-tropic additives in organic solar cells may increase their power efficiency (Karak et al., 2015). It was previously reported that roots can reduce the molecular electron acceptors FeCN and 2,6-dichlorophenol indophenol (DCPIP) (Pardha-Saradhi et al., 2014).

In this work, we follow the ability of green onion roots to reduce Cytochrome C by absorption measurements. We show for the first time that electricity can be harvested directly from green onion roots by associating them with the anode in an electrochemical cell. This electricity generation is enhanced upon the illumination of the onions’ leaves. We apply cyclic voltammetry and 2D fluorescence to detect the electron mediators NADH and NADPH (NAD(P)H) in the external matrix of the roots, suggesting that like in photosynthetic organisms based BPECs, NAD(P)H perform MET between the roots and the anode. Finally, we design a supercapacitor by laser ablation of graphene and show that it can be charged directly charged by the green onion’s roots.

## 2. Materials and methods

### Materials

All chemicals were purchased from Merck unless mentioned otherwise.

### Absorption measurements

O.D. measurements of the cells were done at 750 nm (Nanodrop 2000 UV-Vis spectrophotometer, Thermo Fisher Scientific).

### 2D – FM measurements

The 2D-FM was conducted using a Fluorolog 3 fluorimeter (Horiba) with excitation and emission slits bands of 4 nm. Evaluation of NAD(P)H concentrations was calculated by the construction of an NADH calibration curve based on increasing concentrations measured at (λ_Ex_ = 350 nm, λ_Em_ = 450 nm).

### Chronoamperometry measurements

CA measurements were conducted using a system designed based on the description in the previous work of Shlosberg et al.(Shlosberg et al., 2022b) CA measurements were performed in 4.5 cm^3^ rectangular transparent glass vessels. Illumination was performed using 2 LEDs that were placed vertically to illuminate the onion’s leaves. The distance between the LEDs and the leaves was ∼10 cm. All measurements were performed in a two-electrode mode without the application of electrical potential on the working electrode (WE). A stainless-steel clip was applied as the WE and a platinum wire as the counter electrode (CE). In all measurements, the current density was calculated based on the surface area of the WE that was totally covered by roots of 0.08 cm^2^.

### Chronopotentiometry measurements

Chronopotentiometry was carried out using a Palmsens4 potentiostat. The potentiostat was connected to the supercapacitor in parallel with a connection to a roots fuel cell. The measurements were conducted under real sunlight whose intensity was measured by a light meter at the maximal height of the green onion leaves, measuring a sunlight intensity of about 150 W/m^2^.

### Cyclic voltammetry measurements

Cyclic voltammetry measurements were conducted using a palmsens4 potentiostat connected to screen-printed electrodes (Basi), with a graphite working electrode (1 mm in diameter), a graphite counter electrode, and a silver coated silver chloride reference electrode. Green onion roots were incubated in 0.1 M NaCl for 10 min in the dark or under white light illumination. Right after the incubation, a drop of 60 μL NaCl solution was placed on the screen-printed electrodes, and the CV was measured in the range of -1 – 1 V, with a scan rate of 0.1 V/s.

### Supercapacitor fabrication

Supercapacitors were fabricated using a previously published method.(El-Kady and Kaner, 2013) Briefly, graphite oxide (GO) was synthesized using a modified Hummer’s method and was made into an aqueous dispersion of 2.7 mg mL^-1^. A sheet of polyethylene terephthalate (PET) plastic film was cut and glued onto a LightScribe disc. 16 mL of the stock GO dispersion were then drop-cast on the PET film. After drying overnight under ambient conditions, the prepared GO on PET film was laser-scribed with the desired interdigitated pattern using a LightScribe DVD burner. Subsequently, the edges of the supercapacitor were painted with conductive silver paint, and copper tape was applied on top to act as a current collector. Finally, polyimide Kapton was used to cover the supercapacitor leaving only the interdigitated electrodes exposed.

## 3. Results and Discussion

### 3.1. Onion roots can reduce the electron acceptor Cytochrome C

We first wished to quantitate the ability of green onion roots to perform external electron transport (EET). To do that, we incubated about 10 roots of green onion in a solution of 30 μM Cytochrome C (Cyt) for 10 min. The absorption spectra of the pure Cyt solution and the roots were measured at 1 min time intervals. The obtained spectra showed an increase in the peak with a maxima at λ = 550 nm that correlated with the increasing incubation time (**Fig. 1a**). This intensity peak derives from the reduction of Cyt.(Larom et al., 2015) However, the increase in the incubation time has also elevated the baseline intensity of the whole measured spectrum λ = 475 – 550 nm, probably because of the secretion of molecules that also absorb in this wavelength range. Therefore, to make a more accurate quantification, the Cyt reduction was calculated by subtracting the intensity at the maxima of the peaks (λ = 550 nm) from their minima (λ = 540 nm) (**Fig. 1b**). This analysis showed that a maximal reduction of Cyt (that representing EET of 0.15 μmol electrons) was obtained after 8 min.

**Fig. 1.**
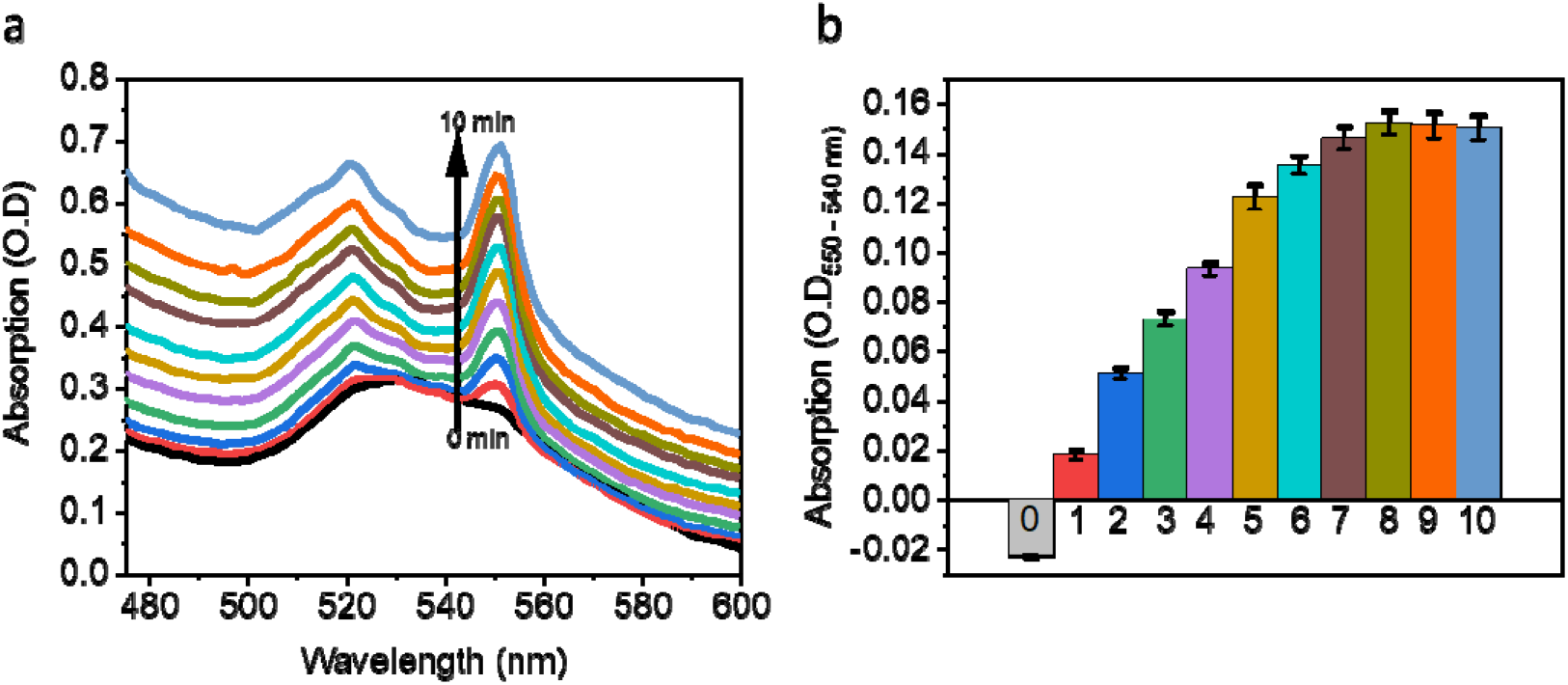
Onion roots reduce the electron acceptor Cytochrome C. Ten green onion roots were incubated for 10 min with Cyt. The absorption spectra were measured before the incubation and then every 1 min. **a** Absorption spectra of the Cyt solution as a function of incubation time. The black arrow represents the time increase that correlates with the intensity increase at λ = 550 nm. **b** The intensities at λ = 550 nm were subtracted by the intensities at λ = 540 to estimate the net absorption increase that derives from Cyt C reduction and is not dependent on the absorption of additional components in the solution. The error bars represent the standard deviation over 3 independent repetitions.

### 3.2. NAD P H is released from roots

Previous studies about fuel cells based on various cells and organisms such as photosynthetic organisms,(Shlosberg et al., 2022d) marine animals, (Shlosberg et al., 2022a) showed that part of the electrical current production derives from the secretion of the redox-active molecules NADH or NADPH (NAD(P)H). In the case of plants, the secretion of NADPH was influenced by its formation in the photosynthesis pathway and was therefore enhanced upon illumination. We postulated that the same molecules may be secreted from roots to reduce external acceptors (Shlosberg et al., 2022c). To elucidate this, green onion roots were incubated in 0.1 M NaCl solution for 10 min in the dark and in high light. The plant (with the roots) was removed, and 2D-fluorescence maps (2D-FM) (λ_Ex_ = 250 - 500 nm, λ_Em_= 280 – 800 nm) were measured (**Fig. 2a**,**b**). The spectral fingerprints of NAD(P)H (λ_Ex max_ = ∼350 nm, λ_Em max_= ∼450 nm), and the amino acids tryptophan and tyrosine (λ_Ex max_ = ∼280 nm, λ_Em max_ = ∼350 nm) were identified based on previous studies (Shlosberg et al., 2021a, 2020). Interestingly, as previously reported for photosynthetic organisms, (Shlosberg et al., 2022d) the intensity of the peaks of NAD(P)H and the amino acids were higher after the illumination of the onion’s leaves. We suggest that NADPH that is formed by photosynthesis may be transported from the leaves to the roots that release it to donate electrons at the anode. To evaluate the NAD(P)H concentrations in the external solution, a calibration curve was prepared by measuring the fluorescence intensity (λ _Ex_= 350 nm, λ _Em_= 450 nm) with increasing NADH concentrations. The NAD(P)H concentrations were determined to be ∼0.15 and 0.27 μM in dark and light, respectively (**Fig. 2c**) (that is equivalent to 15 and 27 nmol). Since unknown reducing molecules rather than NAD(P)H may be secreted from the roots, we wished to estimate the contribution of NAD(P)H to the total reduction of Cyt reduction (**Fig. 1**). To evaluate this, the number of secreted NAD(P)H molecules in the dark was divided by the number of reduced Cyt molecules over 10 min (each NAD(P)H or Cyt molecule can donate/accept 1 electron). The calculation shows that NAD(P)H can reduce about 10% of the Cyt molecules. Based on these results, we suggest that the roots secrete additional reducing molecules rather than just NAD(P)H. However, it is possible that more than 10% of Cyt derives from NAD(P)H reduction if some of the NAD(P)^+^ or NAD(P)H molecules are being up taken up by the roots. Such a cyclic electron transport mechanism was previously reported for various cyanobacterial species(Shlosberg et al., 2020) and microalgae(Shlosberg et al., 2021b) in which NAD(P)H was the major electron mediator. Roots secrete many kinds of components such as minerals, proteins, carbohydrates, and vitamins (Greisenegger, J. K.; Vorbuchner, 1920; Gvamichava, 1966). We postulated that some of the reported vitamins that are capable of charge transfer such as ascorbic acid, thiamine, and nicotinic acid, may also play a role in the reduction of Cyt. To further study the released redox active species, we applied cyclic voltammetry measurements on an 0.1 M NaCl solution after incubation for 10 min with the roots in the dark and in the light (**Fig. 2d**). A small peak was obtained around 0.5 V in the dark, which may originate from the presence of ascorbic acid (Shlosberg et al., 2023b). After illumination, a bigger peak appeared around 0.6 V (that may have masked the small peak at 0.5 V) which may correspond to NADPH (Wang et al., 2021). The diagonal trend of the obtained voltammogram indicates the presence of non-conductive organic materials in the measured solution whose release from the roots is in agreement with previous studies (Moqsud et al., 2017; Strik et al., 2008).

**Fig. 2.**
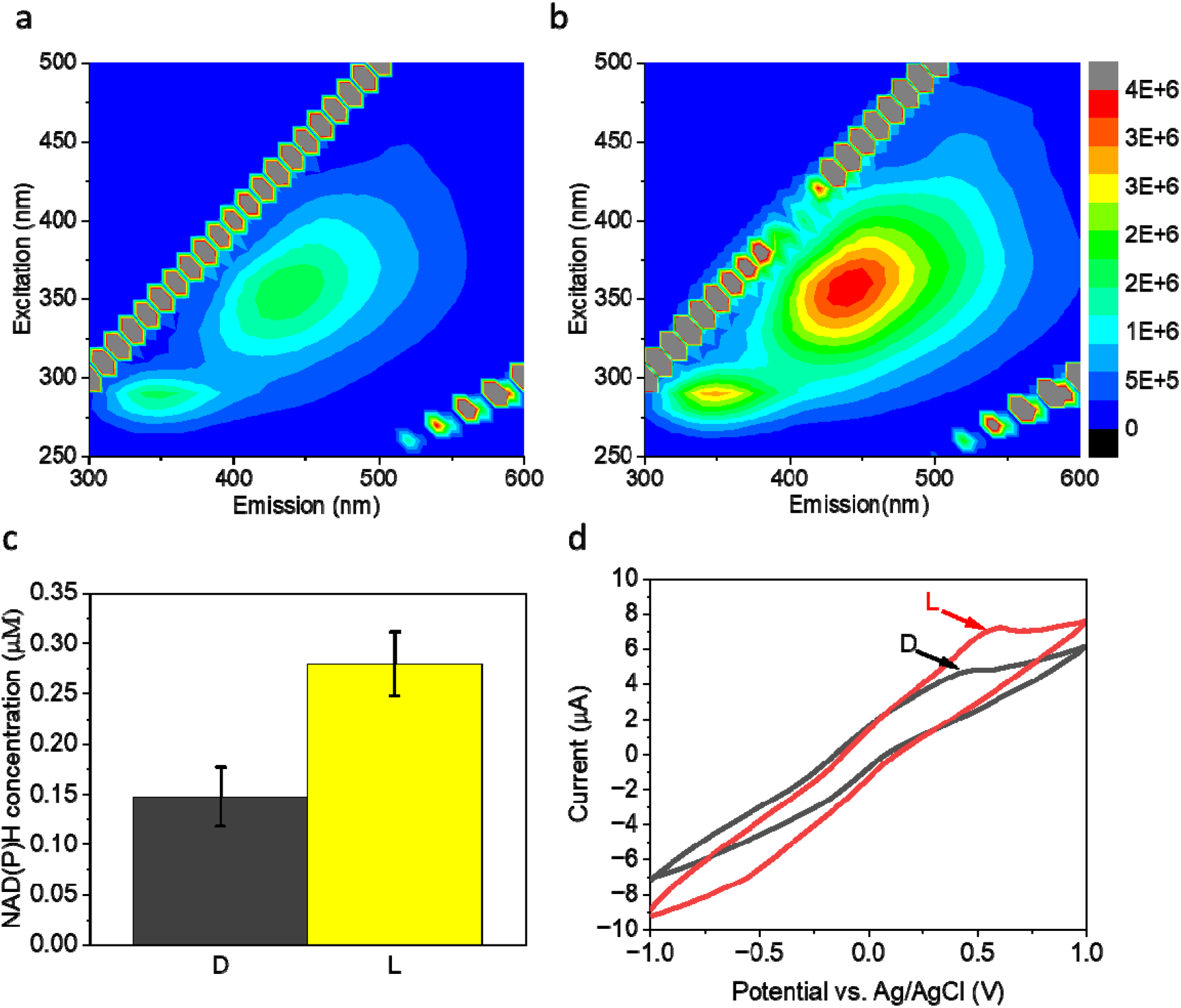
NAD(P)H is secreted from roots. Ten green onion roots were incubated in 0.1 M NaCl solution for 10 min with or without the illumination of the onion’s leaves. Following the incubation, the 2D-FM and cyclic voltammetry of the solutions were measured. **a** 2D-FM after incubation in the dark. **b** 2D-FM after incubation in light. **c** NAD(P)H concentrations were calculated for the solutions after incubation in the dark (D) and in the light (L). The error bars represent the standard deviation over 3 independent repetitions. **c** Cyclic voltammograms after incubation with roots in the dark (D, black) and in the light (L, red).

### 3.3. Harvesting bio-electricity from roots

Based on our results (**Fig. 1**), and previous works that reported the ability of roots to reduce electron acceptors in solution, (Pardha-Saradhi et al., 2014) we thought that the roots could produce an electric current in a bio-fuel cell. For this purpose, a new setup was designed (**Fig. 3a-c**). The setup consists of a rectangular glass vessel with width, length, and height of 4.5 cm. 50 mL NaCl solutions with increasing concentrations of 0.1 – 0.5 M were used as electrolytes. The bottom of the green onion and its roots were placed in the solution. 0.2 cm of the edges of 5 roots were attached to stainless-steel anode clips, to cover the entire anodic area of 0.04 cm^2^. A platinum wire was used as a cathode. The illumination of the leaves was conducted by 2 white LEDs that were placed on a stand ∼10 cm from the leaves. A similar setup that utilizes a stainless steel clip as an anode was previously reported for seaweeds and plant leaves based BPECs (Shlosberg et al., 2022b, 2022c). To optimize the NaCl concentration, Chronoamperometry (CA) of increasing NaCl concentrations (100 – 500 mM) was measured using a 2-electrode mode without the application of an electrical potential bias (bias-free) for 24 h in the dark. The maximal current density that was obtained for the NaCl concentrations: 0.1, 0.2, 0.3, 0.4, and 0.5 M were ∼4.5, 8, 20, 35, and 40 μA/cm^2^, respectively (**Fig. S1**). This electrolyte concentration of 500 mM was also previously determined to be optimal in seaweed and plant leave-based BPECs (Shlosberg et al., 2022b, 2022c).

**Fig. 3.**
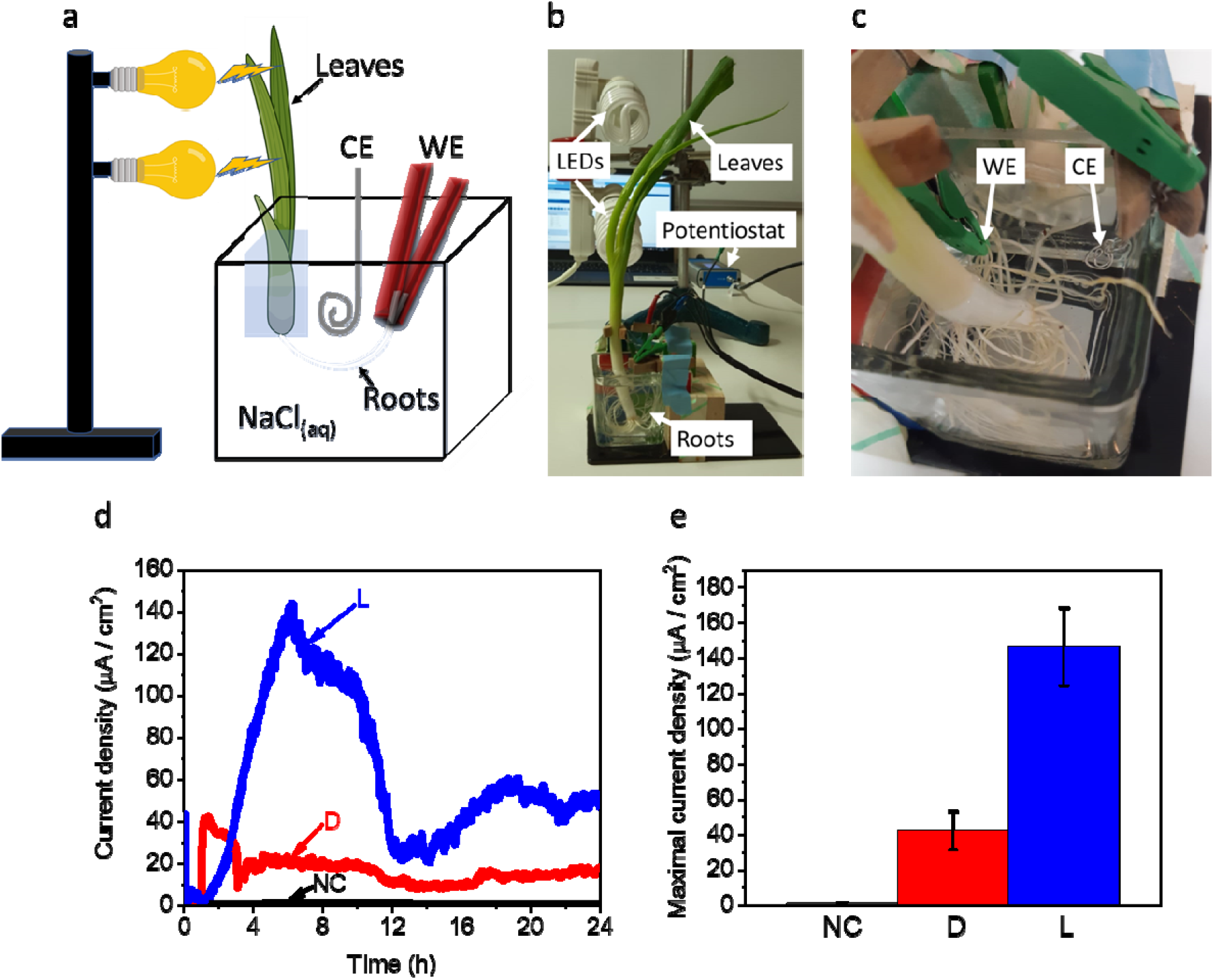
Harvesting bioelectricity from roots. Bias-free CA measurements of green onion roots in a bio-fuel cell in the dark and in light. **a** A schematic illustration of the system: the bottom of a green onion was placed inside the NaCl solution. The edge of the roots was attached to stainless-steel clips (red) that were used as the working electrode (WE), and a platinum wire (gray) was used as a counter electrode (CE). The leaves of the onion were illuminated by white LEDs (100 W) that were placed on a stand at a distance of 10 cm from the leaves. **b** A picture of the measurement system: white labels with arrows mark the roots, leaves, potentiostat, and LEDs. **c** An enlargement of the BEC system: white labels with arrows mark the WE, and CE. **d** CA measurements of green onion roots were conducted over 24 h in the dark (D), in the light (L), and with roots that were not connected to the anode in the light (NC). **e** Maximal current over 24 h. The error bars represent the standard deviation over 3 independent measurements.

Next, we wished to assess whether different pH values affect the current production. For this purpose, the CA of roots was measured using 500 mM NaCl dissolved in MES buffer with pH values of 5 - 8. The change in the pH in this range did not have any significant influence on the current production (**Fig. S2**). Previous studies on BPECs that were based on photosynthetic micro, (Shlosberg et al., 2021b, 2020) and macro-organisms (Shlosberg et al., 2022b, 2022c) suggested that although the mechanism of the electron transfer as conducted by diffusive electron mediators, the organism must be in a close approximation with the anode to supply a sufficient concentration of the mediator. We wished to explore if roots also must be attached to the anode to produce current and whether illumination of the leaves can influence the current production of the roots. Bias-free CA measurements of the onion roots were conducted in 0.5 M NaCl for 24 h in the dark, in the light and without a direct association of the roots with the anode (the roots were inside the electrolyte solution but not touching the anode). No significant current was obtained for the roots without a direct association to the anode. The maximal current densities of the associated roots in the dark and in the light were ~40 and 140 μA/cm^2^, respectively (**Fig. 3d**,**e**). We suggest that the dark current derives from the secretion of NADH, and vitamins such as ascorbic acid, thiamine, nicotinic acid, while the current enhancement in light derives from the trafficking of NADPH molecules that are formed in photosynthesis towards the roots, and outside to donate electrons at the anode.

### 3.4. Charging a supercapacitor by a roots fuel cell

Electricity production may be used by connecting the biofuel cell directly to the electric grid. However, in many cases, the production is conducted far from the grid or generates more electricity than is needed at a specific time point. To improve energy management, the energy may be stored and used in a time of need. To demonstrate the ability to store energy that is produced by roots bio-fuel cells, we wished to charge a supercapacitor. We fabricated a supercapacitor that consists of graphene/graphene oxide (**Fig. 4a**) by laser ablation as reported in our previous work.(Shlosberg et al., n.d.) We designed a system that consists of 2 beakers with a 50 mL solution of 0.5 M NaCl. The first beaker consisted of green onion stainless steel anode clips and a platinum wire as a cathode. Five roots were connected to the anode. Connective leads were wired from the anode and cathode to the graphene/graphene oxide supercapacitor that was placed inside 0.5 M NaCl solution in the second beaker. A potentiostat was connected to the supercapacitor in parallel with the bio-fuel cell (**Fig. 4b**). Chronopotentiometry (CP) was measured under real sunlight illumination (150 W/m^2^) for 10 min. A NaCl solution without the roots was applied as a control experiment. A potential increase was obtained only with the onion roots reaching a maximal potential of 4.5 V after 10 min (**Fig. 4c**,**d**).

**Fig. 4.**
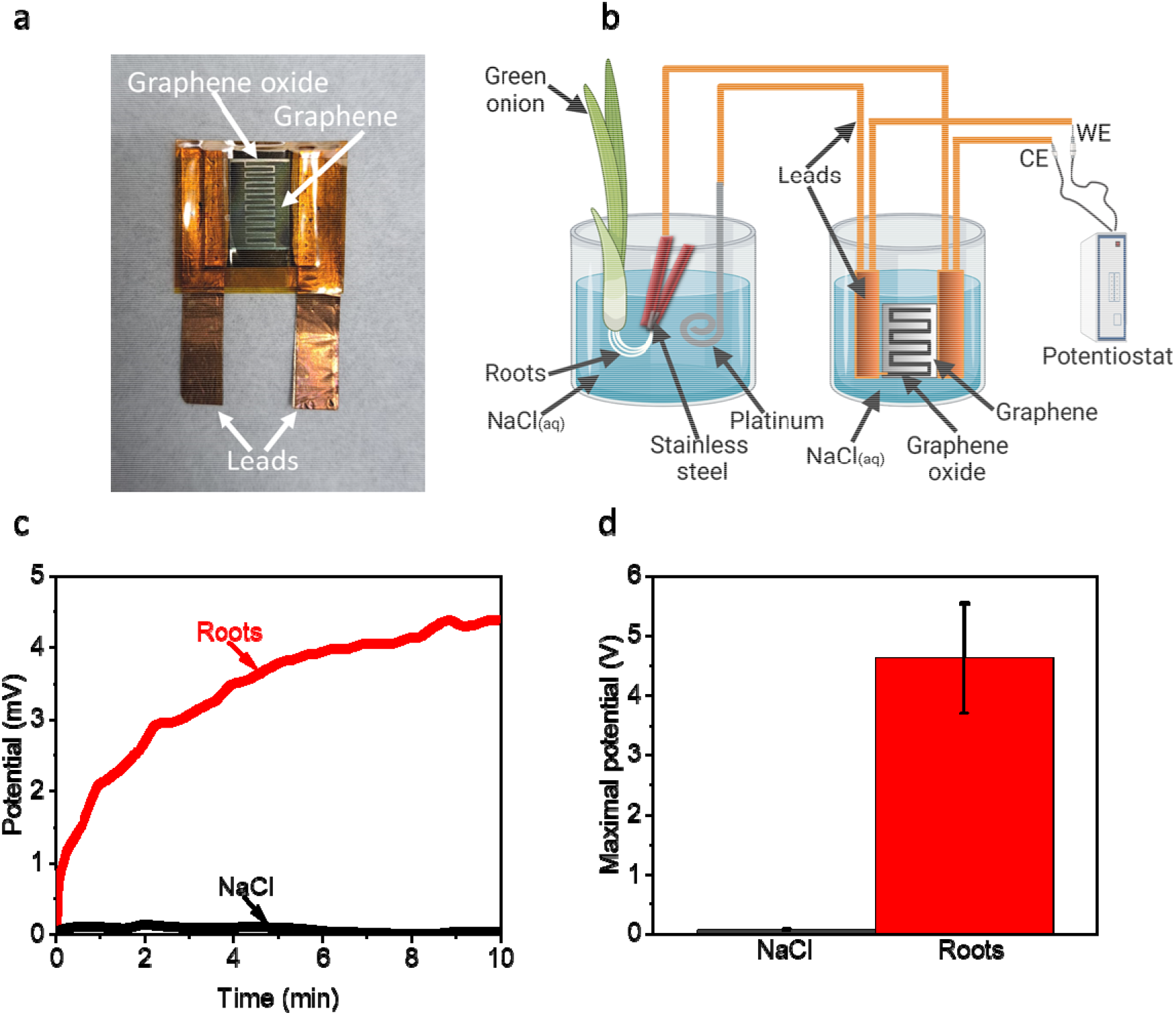
Charging a supercapacitor by a roots fuel cell. A biofuel cell based on green onion roots was applied to charge a graphite oxide/graphene supercapacitor. **a** A photo of the supercapacitor. **b** A schematic description of the system. The system consists of 2 beakers with 0.5 M NaCl solution. The first beaker is the bio-fuel cell which consists of the bottom of a green onion with 5 roots connected to a stainless-steel anode and a platinum cathode. The anode and the cathode were wired to a graphene/graphene oxide-based supercapacitor placed inside the second beaker. The supercapacitor was also connected in parallel to the working electrode (WE) and counter electrode (CE) leads to a potentiostat to measure its charging. The experiment was conducted under real sunlight conditions (∼150 W/m2). **c** Chronopotentiometry measurements with and without the green onion roots. **d** Maximal potential with and without roots over 10 min. The error bars represent the standard deviation over 3 independent repetitions.

### 3.5. Proposing a mechanism for the electrical current production in root fuel cells

Previous studies showed that the metabolic pathways of photosynthesis and respiration are connected, while the NADH molecules that are formed in the TCA cycle can reduce the plastoquinone molecules that proceed into multistep redox reactions that end in the formation of NADPH (Saper et al., 2018). In our work, we showed that NADH and NADPH are being released from the roots, while their excretion is enhanced upon illumination. Based on the previous work and ours, we propose a mechanism for electron transfer (**Fig. 5**). NADH and other unknown redox-active biomolecules such as vitamins can be released from the roots to donate electrons at the anode. Upon solar illumination, NADPH is formed in the photosynthetic pathway while some of it moves from the leaves to the roots and is released to donate electrons at the anode. The oxidized forms NAD^+^ and NADP^+^ may be taken up by the roots and be reduced inside the onion.

**Fig. 5.**
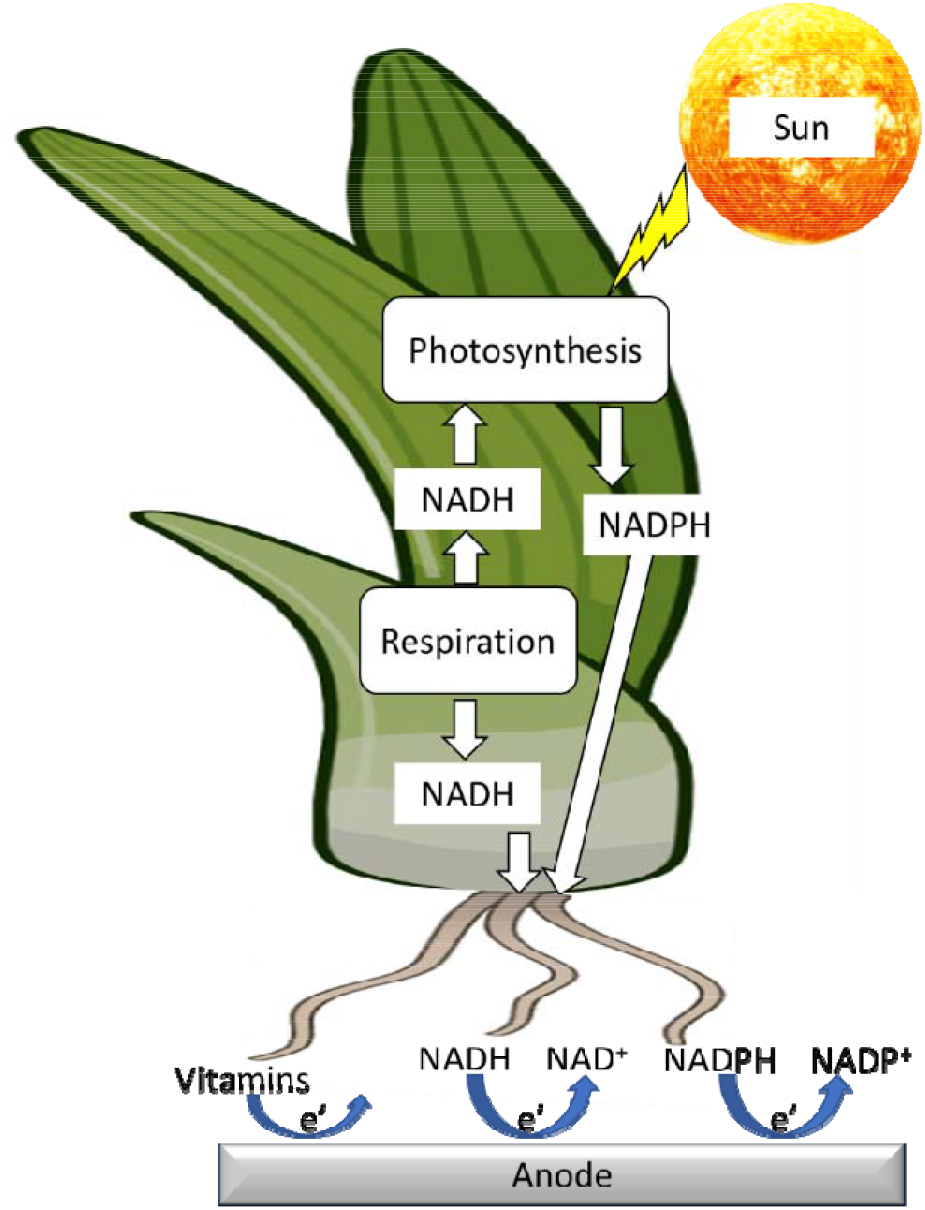
Proposed mechanism for the electrical current production in root fuel cells. A schematic illustration of the suggested electron transport mechanism. NADH molecules that are formed in the respiration pathway can migrate toward the roots or continue to the photosynthesis pathway that produces NADPH, which can also migrate toward the roots. NADH, NADPH, and other unknown redox-active molecules can exit the roots to donate electrons at the anode to produce current. The yellow lightening bolt represents illumination from the sun. White arrows represent the movement of NADH and NADPH between the metabolic pathways of respiration and photosynthesis and their physical movement toward the roots. The round blue arrows represent the reduction of the anode by NADH, NADPH, and additional unknown redox-active molecules.

## 4. Conclusions

In this work, we show that the roots of green onions can release molecules that can reduce the electron acceptor Cyt. We apply 2D-fluorescence and cyclic voltammetry measurements to identify the secretion of the electron mediators NADH and NADPH from the roots. We show for the first time a bio-fuel cell that can produce bias-free electricity directly from intact roots for more than 24 h. This current production is significantly enhanced by the illumination of the onion’s leaves. Finally, we show the ability of the roots fuel cell to store energy by charging a supercapacitor. The ability to produce electrical current from roots especially in hydroponic cultures may pave the way toward the establishment of novel renewable energy technologies that supply the energy demand of green houses facilities. Such technologies can be based on a circular economy model in which agricultural crops will be used for both food and energy generation and storage.

## Acknowledgments

Yaniv Shlosberg is supported by the Otis Williams Fellowship. Some of the results reported in this work were obtained using central facilities at the UCSB Materials Research Laboratory. We thank Jaya Nolt for her technical support. R.B.K. thanks the Dr. Myung Ki Hong Endowed Chair in Materials Innovation at UCLA. Some of the figures were prepared using Biorender.com

## Author contributions

Y.S. conceived the idea. Y.S. and A.H. designed the experiments. Y.S. and A.H. performed the main experiments. Y.S., A.H, T.N.T., and R.B.K. wrote the paper. T.N.T and R.B.K. supervised the entire research project.

## Supporting Information for

**Fig. S1.**
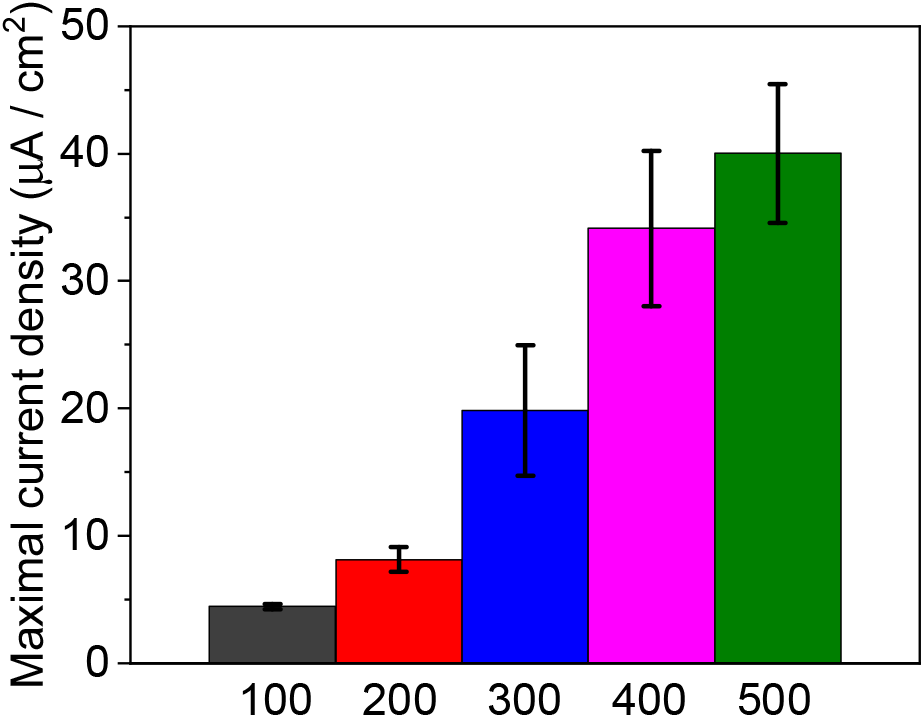
Electric current production from roots at increasing electrolyte salinity. CA of onion roots measured in water solutions with various increasing salinities of 100, 200, 300, 400, and 500 mM NaCl. The error bars represent the standard deviation over 3 independent repetitions.

**Fig. S2.**
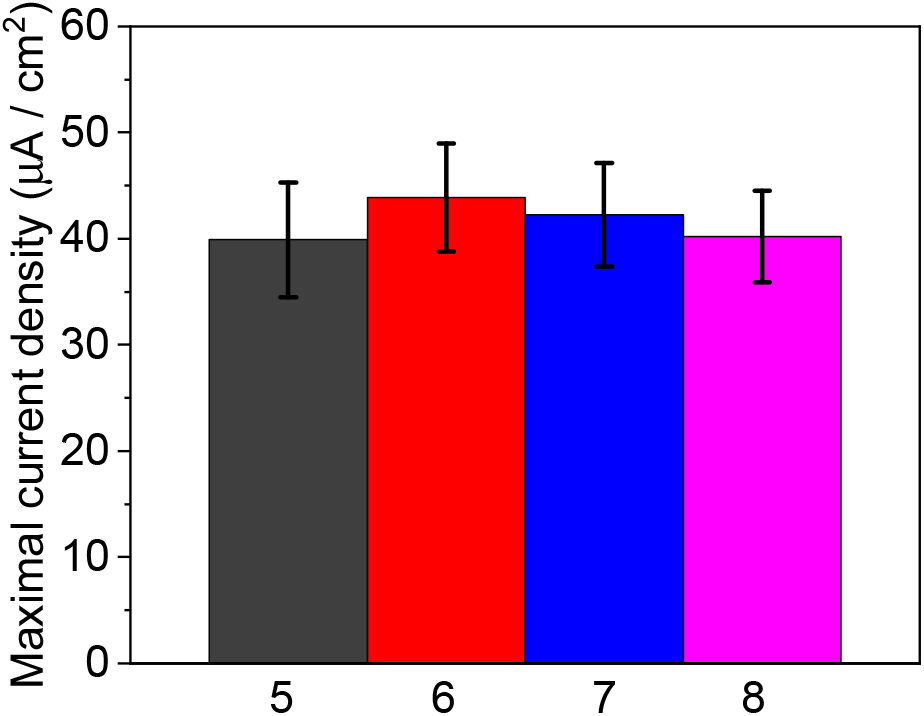
Electric current production from roots at increasing pH values. CA of onion roots measured in a solution of 500 mM NaCl + MES buffer with increasing pH values of 5, 6, 7, and 8. The error bars represent the standard deviation over 3 independent repetitions.

